# AMP-kinase mediates adaptation of glioblastoma cells to conditions of the tumour microenvironment

**DOI:** 10.1101/2022.03.25.485624

**Authors:** Nadja I. Lorenz, Benedikt Sauer, Pia S. Zeiner, Maja I. Strecker, Anna-Luisa Luger, Dorothea Schulte, Michel Mittelbronn, Tijna Alekseeva, Lisa Sevenich, Patrick N. Harter, Christian Münch, Joachim P. Steinbach, Michael W. Ronellenfitsch

## Abstract

AMP-activated protein kinase (AMPK) is a central cellular energy sensor that regulates metabolic activity. We hypothesised that in glioblastoma (GB), AMPK plays a pivotal role in balancing metabolism under conditions of the tumour microenvironment, which is characterised by fluctuating and often low nutrient and oxygen availability. Impairment of this network could thus interfere with tumour progression.

AMPK activity was modulated genetically by CRISPR/Cas9-based double knockout (DKO) of the catalytic α1 and α2 subunits in human GB cells and effects were confirmed by pharmacological AMPK inhibition using BAY3827 and an inactive control compound in primary GB cell lines.

We found that metabolic adaptation of GB cells under energy stress conditions (hypoxia, glucose deprivation) was dependent on AMPK and accordingly that, AMPK DKO cells were more vulnerable to glucose-deprivation or inhibition of glycolysis and sensitised to hypoxia-induced cell death. This effect was rescued by reexpression of the AMPK α2 subunit. Similar results were observed using the selective pharmacological AMPK inhibitor BAY3827. Mitochondrial biogenesis was regulated AMPK-dependently with a reduced mitochondrial mass and mitochondrial membrane potential in AMPK DKO GB cells. *In vivo*, AMPK DKO GB cells showed impaired tumour growth and tumour formation in CAM assays as well as in an orthotopic glioma mouse model.

Our study highlights the importance of AMPK for GB cell adaptation towards energy depletion and emphasises the role of AMPK for tumour formation *in vivo*. Moreover, we identified mitochondria as central downstream effectors of AMPK signalling. The development of AMPK inhibitors could open opportunities for the treatment of hypoxic tumours.

## Introduction

Glioblastoma (GB) is a heterogeneous diffuse glial tumour with distinct histological features including necrosis and neoangiogenesis that mirror nutrient deprivation and hypoxia in the tumour microenvironment, which has been shown to be fluctuating. The resulting cycles of hypoxia and starvation are thought to drive progression of solid tumours ^1^. Standard therapy comprises surgery followed by radiotherapy and alkylating chemotherapy with temozolomide in patients in sufficient clinical condition ^2^. Recent clinical development has failed to identify additional drugs for the narrow arsenal of GB treatment options, thus novel therapeutic strategies are urgently needed.

Altered metabolism has been recognized as a hallmark of cancer and might expose a targetable Achilles’ heel ^3^. Glucose is considered the major energy source for cancer cells and oxidation *via* glycolysis and subsequent oxidative metabolism generates energy by increasing adenosine triphosphate (ATP) levels. While human physiology aims at maintaining a steady state serum spectrum of nutrients and oxygen, cells are additionally equipped with an intrinsic machinery to adjust metabolism based on energy supply ^4^. At the core of this mechanism is 5′-adenosine monophosphate (AMP)-activated protein kinase (AMPK) which is activated by an increase of the AMP/ATP ratio ^5^ as well as glucose deprivation *via* the glycolytic enzyme aldolase ^6,7^ and orchestrates adaptation of metabolism during nutrient starvation conditions to promote cell survival. The heterotrimeric protein AMPK is composed of a catalytic α subunit that contains the phosphorylation site Thr172 which is essential for activation and the regulatory β and γ subunits ^8^. Notably, several isoforms of these proteins encoded by distinct genes exist in mammalian cells. During energy stress conditions AMP or ADP binding to the γ subunit of AMPK triggers phosphorylation of Thr172 of the α subunit by upstream kinases, mostly by the liver kinase B1 (LKB1)^9^. To maintain energy homeostasis in states of low external energy supply, AMPK directs metabolism towards catabolism e.g. by activation of fatty acid oxidation and glycolysis for ATP production ^5^. AMPK phosphorylation targets include acetyl-CoA carboxylase (ACC), which catalyses the first step of fatty acid synthesis and is inhibited by Ser79 phosphorylation. This phosphorylation is frequently used as a surrogate marker for AMPK activity ^10,11^. *Via* phosphorylation of its targets tuberin (TSC2) and Raptor, AMPK also indirectly inhibits mammalian target of rapamycin complex 1 (mTORC1) signalling which integrates signalling from growth factor receptors with the metabolic state of the cell to regulate diverse targets involved in translation, cell growth and autophagy ^12^.

Besides its role in metabolic programming to counteract starvation conditions, AMPK regulates mitochondrial dynamics including fusion and fission states and homeostasis as well as mitochondrial quality control by regulation of mitophagy ^13,14^. AMPK also regulates energy homeostasis under glucose starvation conditions *via* an AMPK-p38MAPK axis that induces the expression of peroxisome proliferator-activated receptor gamma coactivator 1-alpha (PGC-1α) in MEFs and H1299-EV cells. In consequence, an enhanced mitochondrial activity and mitochondrial biogenesis ensures cellular survival under glucose-limiting conditions ^15^.

It appears plausible that AMPK exerts important functions for cellular stress adaptation in tumours. However, there are conflicting results with regard to pro- and anti-tumour effects for different types and stages of cancer ^16–18^. The fact that lung cancers frequently have LKB1 mutations with potentially reduced levels of AMPK activity ^19^ indicates a tumour suppressive role of AMPK. In contrast, recently, a chronic activation of AMPK by oncogenic stress in GB cells has been reported to regulate hypoxia-inducible factor 1α and glycolysis *via* phosphorylation of the transcription factor CREB1 to enhance tumour growth and GB bioenergetics ^18^. In this context, inhibition of AMPK by gene suppression of the β1 regulatory subunit led to reduced tumour growth *in vitro* and *in vivo* together with reduced glycolysis and oxidative phosphorylation ^18^. While these results comply with an oncogenic role of AMPK in GB, AMPK activation has also been suggested as a therapeutic target for activation in GB. So far, all available pharmacologic AMPK inhibitors lack specificity and hence are not ideally suited for investigations on AMPK dependent effects in preclinical studies and clinical studies. For example, the established AMPK inhibitor Compound C has often been used to investigate AMPK-dependent cellular effects in many types of cancer, nevertheless it interferes with several other kinases leading to AMPK-independent effects ^20–22^.

In this experimental study, we investigated the role of the AMPK-mediated stress response in human GB cells. Mimicking the conditions of the tumour microenvironment we deprived GB cells of glucose and oxygen to characterize metabolic effects and to delineate the relevance of potential downstream targets. AMPK inhibition was modelled by using double-knockout cells of the catalytic subunits α1 and αβ. We here report that AMPK activation promotes GB cell survival in the context of both hypoxia and glucose starvation by promoting metabolic adaptation. Furthermore, we found that knockout of the catalytic AMPK subunits resulted in an impaired tumour formation and prolonged survival *in vivo*. Taken together, the results of this study indicate that the use of AMPK inhibitors could be beneficial to improve GB therapy strategies.

## Material and methods

### Reagents, cell lines and culture conditions

LNT-229 cells have been described ^23^. LN-308 cells were a gift of Nicolas de Tribolet (Lausanne, Switzerland); G55T2 cells were provided by Manfred Westphal and Kathrin Lamszus (Hamburg, Germany) ^24^. All cell lines were maintained in cell culture incubators at 37 °C under a 5% CO_2_ atmosphere. Cells were cultured in Dulbecco’s modified eagle medium (DMEM) containing 10% fetal calf serum (FCS) (Thermo Fisher Scientific, Hamburg, Germany), 100 IU/ml penicillin and 100 μg/ml streptomycin (Life Technologies, Karlsruhe, Germany) ^23^.

LNT-229 and LN-308 cells were authenticated by STR analysis (Multiplexion, Heidelberg, Germany). The STR profile of LNT-229 cells matched with the known profile for LN-229, which only differ in their p53 status ^25^. A STR profile of G55T2 cells has not been deposited in databases yet ^26^.

All reagents, if not specified elsewhere, were purchased from Sigma (Taufkirchen, Germany). BAY974 and BAY3827 were kindly provided by the DCP (Donated Chemical Probes) program ^27^.

### Generation of CRISPR/Cas9 knockout cells

For CRISPR/Cas9 knockout AMPK sgRNA plasmids targeting exon 1 of AMPK α1 and αβ (pX462-hPRKAA1-gRNA, pX462-hPRKAA2-gRNA, #74374-74377) were purchased from Addgene (Watertown, MA, USA). LNT-229, G55T2 and LN-308 cells were transfected with a combination of the gRNA plasmids (0.625 μg each) using Lipofectamine3000 (Thermo Fisher Scientific, Hamburg, Germany) for 6 h. After 3 days, cells were selected with puromycin (2 μg/ml). Single cell clones were analysed for AMPK expression by immunoblotting.

### Primary cell culture

P3NS primary cells were a gift of Simone Niclou (Luxembourg Institute of Health, Luxembourg) and have been characterized elsewhere ^28^. P3NS cells were cultured in Neurobasal A medium supplemented with 1x B27 supplement, 2 % glutamine, 1 U/ml heparin, 20 ng/ml epidermal growth factor (EGF) and 20 ng/ml human recombinant basic fibroblast growth factor (bFGF) (ReliaTech, Wolfenbüttel, Germany).

### Transfection of cells

For expression of wildtype AMPK α2 (*PRKAA2*) in LNT-229 AMPK α1/2 double knockout cells, pcDNA3.1 plasmids with the according constructs were purchased from GenScript (Leiden, The Netherlands). Transfection with Attractene (Qiagen, Venlo, Netherlands) was performed according to the manufacturer’s protocol and the empty pcDNA3.1 plasmid was used as control. For selection medium containing 400 μg/ml G418 was used. The efficacy of transfection was checked in early passage pooled clones.

### Induction of hypoxia

Hypoxia was induced with incubation in GasPak pouches for anaerobic culture (Becton-Dickinson, Heidelberg, Germany) as previously described ^23,29,26^. Experiments in hypoxia (0.1% O_2_) or normoxia (21% O_2_) were performed in serum-free DMEM adjusted to 2 mM glucose.

### Cell density and viability assays

Cell density was assessed by crystal violet (CV) staining as previously described ^30^. For cell viability measurements propidium iodide (PI) uptake was quantified by flow cytometry (FACS) as previously described ^23^. A BD Canto II flow cytometer was used for data acquisition and data analysis was performed with the BD FACS Diva software version 6.1.3. Cell viability analysis by lactate dehydrogenase (LDH) release assay was performed with the Cytotoxicity Detection Kit (LDH) (Roche, Mannheim, Germany) and has also been described ^23,31^.

### Immunoblot analysis

Immunoblot analyses were performed following a standard protocol ^32^ with the following antibodies: pACC (Ser79), pAMPK (Thr172), ACC and AMPKα1/2 (#3661, #2535, #3661, #2532, Cell Signaling Technology, Danvers, MA, USA), AMPKα2 (#18167-1, Proteintech, Rosemont, IL, USA) and actin (#sc-1616, Santa Cruz Biotechnology, Santa Cruz, CA, USA). Secondary anti-rabbit and anti-goat antibodies were purchased from Jackson ImmunoResearch (#111-036-144; West Grove, PA, USA) and Santa Cruz Biotechnology (#sc-2020).

### RNA isolation and quantitative real-time PCR

RNA extraction and cDNA synthesis was performed as described ^32^. Absolute SYBR Green Fluorescein q-PCR Mastermix (ThermoFisher Scientific, Hamburg, Germany) was used for quantitative RT-PCR measurements with the corresponding primers. *18S* and *SDHA* were used as housekeeping genes for normalisation.

### Determination of mitochondrial mass and mitochondrial membrane potential by flow cytometry

Changes in mitochondrial mass and mitochondrial membrane potential were measured by flow cytometry. Briefly, cells were seeded and allowed to attach overnight. Cells were treated as indicated. After washing with PBS, cells were stained with 100 nM MitoTracker Green FM or Mitotracker Red (Thermo Fisher Scientific, Hamburg, Germany) for 20 min. Afterwards cells were harvested, washed and analysed by flow cytometry (BD Canto II).

### *In vivo* experiments

All *in vivo* mouse experiments were approved by the local animal ethics committee (regional board Darmstadt, Germany), four-week-old athymic nude mice (Crl:NU (NCr) Foxn1^nu^) were purchased from Charles River (Sulzfeld, Germany). Animals were fed with standard food and water ad libitum and housed on a 12 h dark/night cycle in the local animal facility. Mice were allowed to acclimate in the local animal facility before start of the experiment. Mice were anesthetised and received metamizol for pain relief. For tumour cell injection, mice were placed into a stereotactic fixation device and 1×10^4^ G55T2 wt or AMPK DKO cells resuspended in 2 μl PBS were injected into the right striatum through a burr hole in the skull using a 10 μl Hamilton syringe. Wounds were closed with Surgibond tissue adhesive. After tumour cell injection, all animals were checked twice daily and sacrificed when neurological symptoms or a loss of more than 20 % of body weight were determined. Brains were isolated and fixed in 4 % PFA for immunohistochemical analysis.

Tumour growth was monitored by MRI. MRI measurements were performed at the Animal Imaging Core Facility after 11, 18 and 25 days using a 7 Tesla small animal MRI (Pharmascan, Bruker) and analysed using ITK Snap software ^33^.

### Statistical analysis

All quantitative data are expressed as mean and standard deviation (SD). P-values were calculated with two-tailed student’s t-tests. Values of p>0.05 were considered as not significant (n.s.), values of p<0.05 and p<0.01 as significant or highly significant.

## Results

### Double knockout of both AMPK catalytic subunits inhibits signalling under energy stress conditions in human GB cells

AMPK is known as a central cellular energy sensor that is necessary to switch to an increased catabolism during cellular energy stress conditions ^8^. To elucidate the effect of AMPK activity for GB cells, LNT-229, LN-308 and G55T2 cells with a double knockout (DKO) of the catalytic subunits α1 and α2 were generated by CRISPR/Cas9 gene editing. While other studies had used GB cells with a knockdown of the regulatory AMPK β subunit ^18^, AMPK α1/α2 DKO cells offer the benefit of lacking catalytic activity and are therefore important tools to investigate AMPK-dependent effects in GB cells. LNT-229, G55T2 and LN-308 DKO cells showed no AMPK specific band in immunoblot analysis (Fig. 1A). Two clones of each cell line were used for further experiments. Concomitant glucose and oxygen starvation, mimicking the conditions of the GB microenvironment with low oxygen and nutrient availability, resulting in an AMPK-mediated phosphorylation of ACC as surrogate marker for AMPK activity, whereas ACC phosphorylation was absent in all tested AMPK DKO GB cell lines (Fig. 1B).

**Fig. 1:**
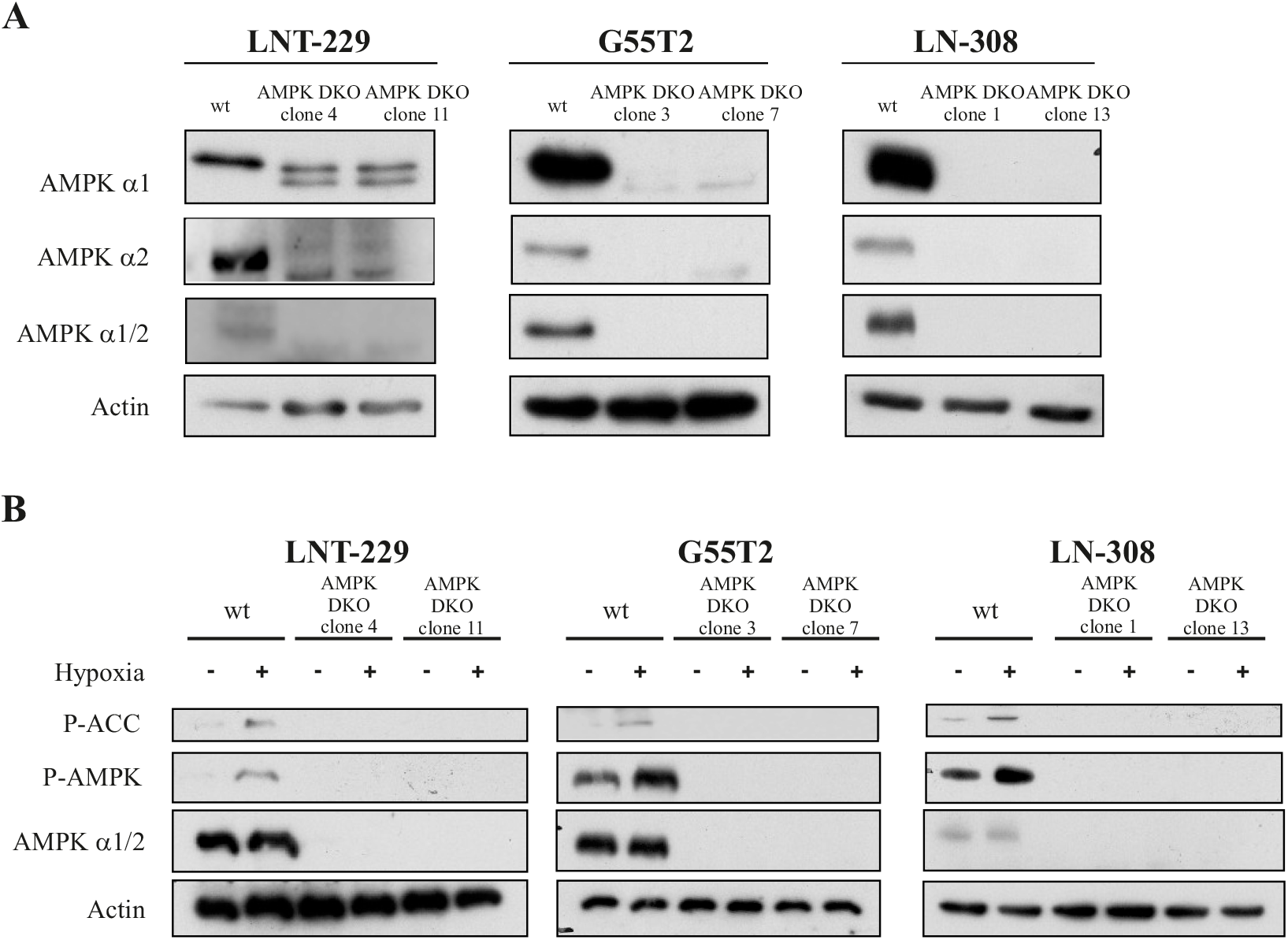
AMPK DKO leads to inhibition of AMPK activation and deregulated metabolic adaptation under hypoxic conditions in human GB cells. (A) Cellular lysates of LNT-229, LN-308 and G55T2 wt and AMPK DKO cells were analysed by immunoblot with antibodies for AMPKα1, AMPKα2, AMPKα1/2 and actin. (B) LNT-229, LN-308 and G55T2 wt or AMPK DKO cells were incubated in serum-free DMEM supplemented with 2 mM glucose for 8 h in normoxia (21 % O_2_) or hypoxia (0.1 % O_2_). Immunoblots with antibodies for P-ACC, P-AMPK, AMPK α1/2 were performed.

### AMPK DKO cells are sensitised to nutrient starvation- and hypoxia-induced cell death

To investigate the effect of defective AMPK signalling in human GB cell lines, we exposed LNT-229, G55T2 and LN-308 AMPK DKO cells to glucose starvation and hypoxia. Cells cultured in glucose-free medium showed a significantly higher rate of cell death compared to corresponding wildtype cells (Fig. 2A). Under hypoxic conditions with reduced glucose availability, the hypersensitivity of LNT-229, G55T2and LN-308 AMPK DKO cells was even more pronounced (Fig. 2B). Besides the genetic model, effects of pharmacological inhibition of AMPK were analysed using the specific chemical probe BAY3827. This potent compound is reported to specifically target the α1 subunit of AMPK ^34^. In LNT-229 cells, treatment with BAY3827 led to reduced P-ACC levels. In contrast, P-AMPK levels increased after BAY3827 treatment (Supplementary Fig. 1A). Under glucose starvation conditions pharmacological AMPK inhibition with BAY3827 also increased cell death compared to vehicle treated cells or cells treated with the inactive, but chemically similar control probe BAY974 (Supplementary Fig. 1B). Moreover, LNT-229 cells treated with BAY3827 were sensitised to hypoxia-induced cell death (Supplementary Fig. 1C). To transfer these observations, P3NS primary GB cells were treated with BAY974 or BAY3827. Comparable to LNT-229 cells, P-ACC levels were reduced after pharmacological AMPK inhibition, whereas P-AMPK levels increased (Fig. 2C). Furthermore, we found that BAY3827 drastically increased cell death when cells were treated with BAY3827 under glucose-free conditions (Fig. 2D). These results indicate an important role of AMPK for the adaptation to conditions of the tumour microenvironment in human GB cells.

**Fig. 2:**
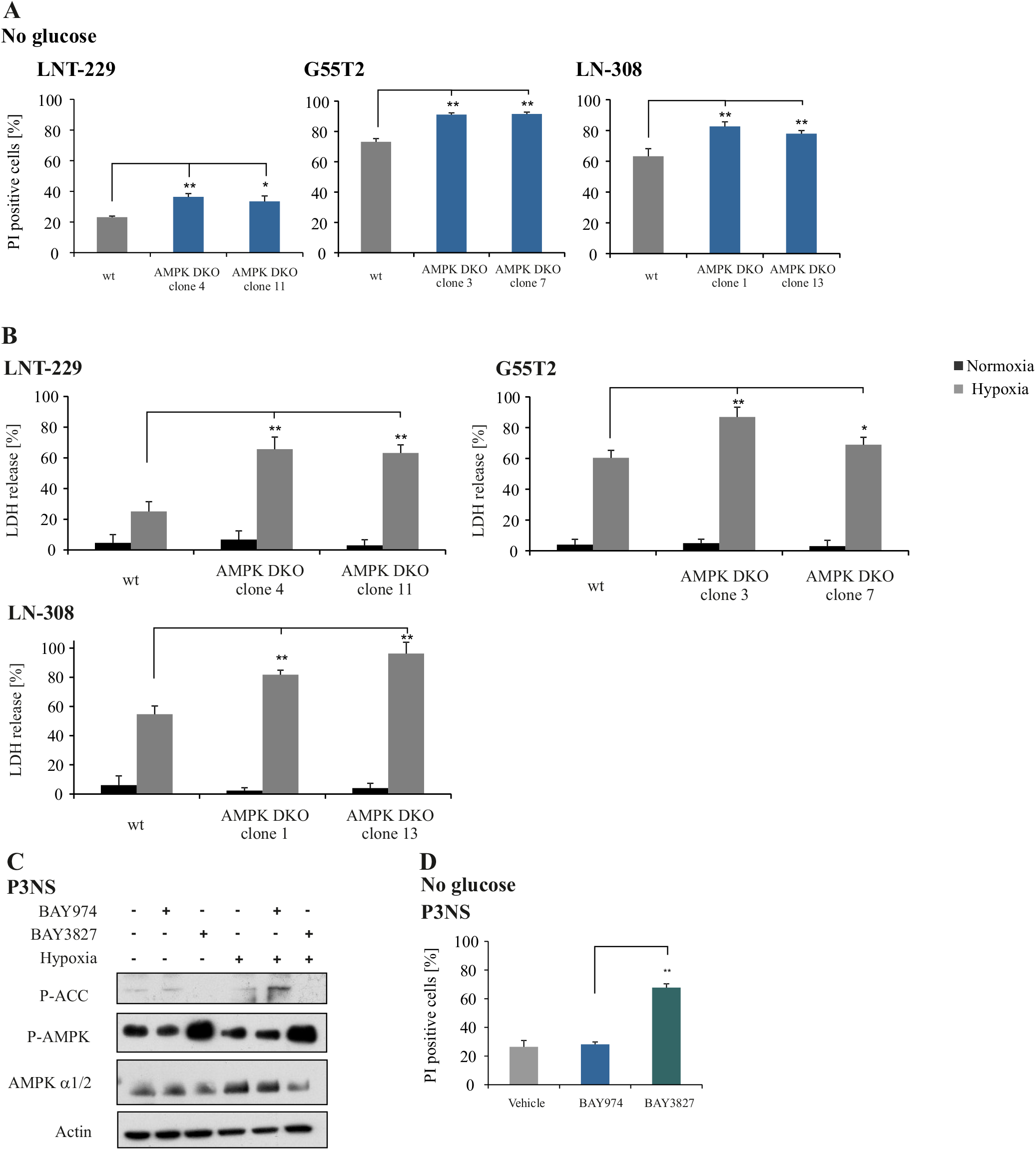
Pharmacological AMPK inhibition and AMPK DKO increase nutrient deprivation- and hypoxia-induced cell death in human GB cells. (A) LNT-229, LN-308 and G55T2 wt and AMPK DKO cells were treated in serum- and glucose-free DMEM for 24 h. Cell death was analysed by PI staining and quantified by flow cytometry (n=3, mean ± SD, **p<0.01, Student’s t-test). (B) Cell death of LNT-229, LN-308 and G55T2 wt and AMPK DKO was analysed by an LDH release assay after incubation of the cells in serum-free DMEM containing 2 mM glucose in normoxia or hypoxia (0.1 % O_2_) (n=4, mean ± SD, *p<0.05, **p<0.01, Student’s t-test). (C) Primary GB cells (P3NS) were treated with vehicle (DMSO), 1 μM BAY974 or 1 μM BAY3827 in serum-free medium containing 2 mM glucose for 8 h in normoxia or hypoxia (0.1 % O_2_). Cellular lysates were analysed by immunoblot with antibodies for P-ACC, P-AMPK, AMPKα1/2 and actin. (D) P3NS cells were treated with vehicle (DMSO), 1 μM BAY974 or 1 μM BAY3827 in serum-free medium without glucose. Cell death was analysed by PI staining and flow cytometry (n=3, mean ± SD, **p<0.01, Student’s t-test).

To confirm AMPK dependency of the observed effects under energy stress conditions, LNT-229 AMPK DKO cells were retransfected with *PRKAA2*, coding for the α2 subunit of AMPK (Fig. 3A). Cells reexpressing the α2 AMPK subunit showed a slight increase in P-ACC levels, which were however still reduced when compared to wildtype cells (Fig. 3B). Nevertheless, *PRKAA2* retransfected LNT-229 AMPK DKO cells showed reduced cell death under nutrient starvation conditions (Fig. 3C). Similarly, retransfected cells were partly protected from hypoxia-induced cell death compared to LNT-229 AMPK DKO cells (Fig. 3 D).

**Fig. 3:**
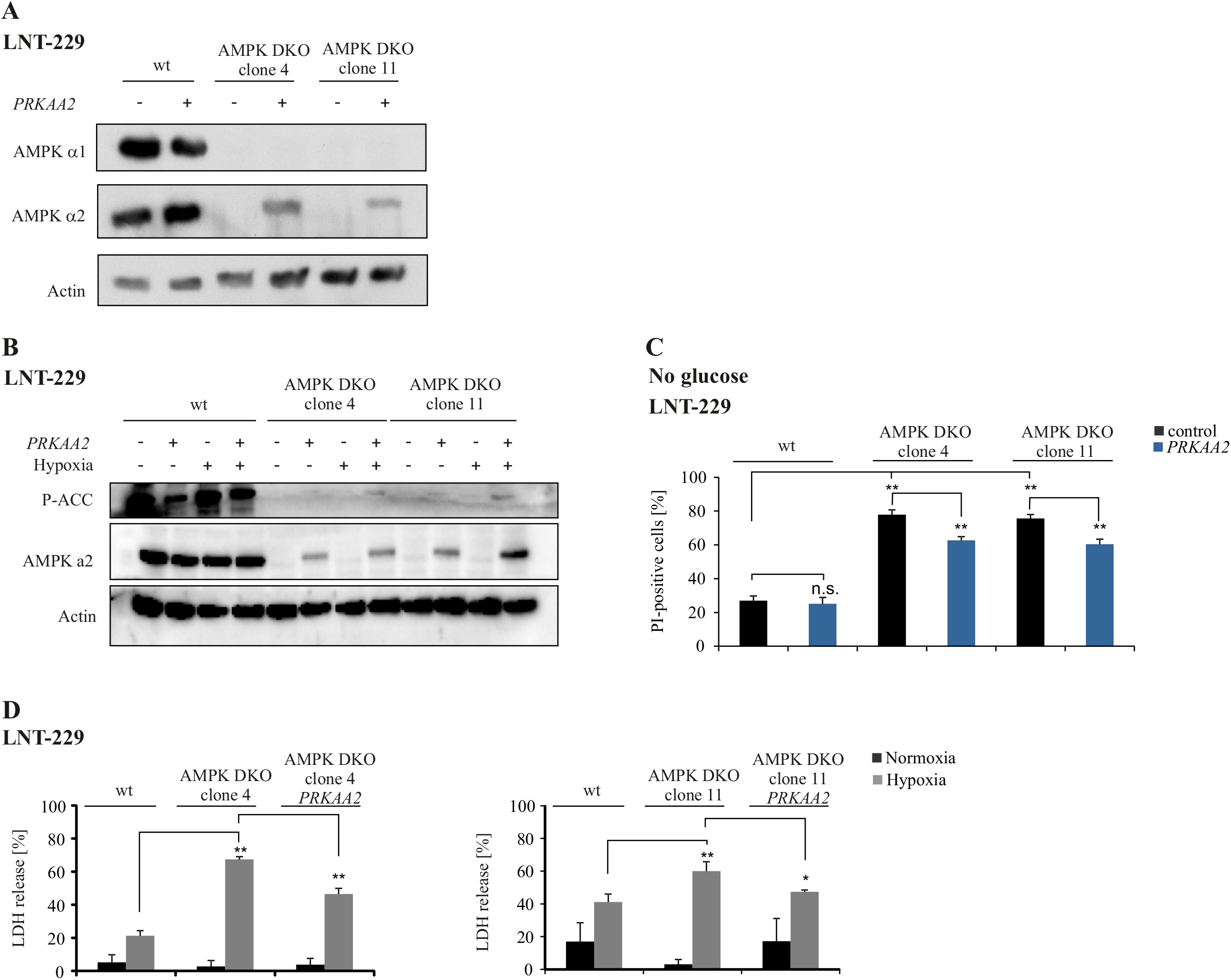
Retransfection of AMPK DKO cells results in decreased cell death under energy starvation conditions. (A) LNT-229 wt and AMPK DKO cells were stably transfected with empty vector (control) or *PRKAA2*. Cellular lysates were analysed by immunoblot with antibodies for AMPK α1, AMPK α2 and actin. (B) Immunoblot analysis of LNT-229 wt and AMPK DKO PRKAA2 lysates treated in serum-free medium with 2 mM glucose for 8 h in normoxia or hypoxia (0.1 % O_2_) was performed with antibodies for P-ACC, AMPK α2 and actin. (C) LNT-229 wt and AMPK DKO PRKAA2 cells were treated in serum-free medium without glucose. Cell death was analysed by PI uptake and flow cytometry (n=3, mean ± SD, n.s. not significant, **p<0.01, Student’s t-test). (D) Cell death of LNT-229 wt and AMPK DKO cells was analysed by LDH release assay after incubation in serum-free medium supplemented with 2 mM glucose in normoxia and hypoxia (0.1 % O_2_) (n=4, mean ± SD, *p<0.05, **p<0.01, Student’s t-test).

### Mitochondrial abundance and activity are dependent on AMPK catalytic functionality

Previous studies have reported that AMPK regulates mitochondrial biogenesis and activity depended on the cellular energy status ^13,14,35^. To investigate whether mitochondrial mass and biogenesis is regulated in an AMPK-dependent manner in human GB cells, LNT-229 and G55T2 wildtype and AMPK DKO cells were analysed for mitochondrial DNA content by qPCR analysis with primers targeting the mtDNA D-loop. In both cell lines AMPK DKO resulted in a reduced mitochondrial DNA content compared to wildtype cells (Fig. 4A). We further analysed mRNA expression levels of several mitochondrial encoded genes as well as mitochondria-associated genes in G55T2 wildtype and AMPK DKO cells (Fig. 4B). Mitochondrial encoded genes (*mtDNA D-loop*, *MT-CYB*, *MT-ND1* and *MT-CO1*) as well as the mitochondria associated gene *ATP5G1* were downregulated in AMPK DKO cells compared to wildtype cells. In line with this observation, mitochondrial abundance (Fig. 4C, left panel) and mitochondrial membrane potential were reduced (Fig. 4C, right panel) which was also confirmed microscopically in LNT-229 wildtype and AMPK DKO cells (Fig. 4D).

**Fig. 4:**
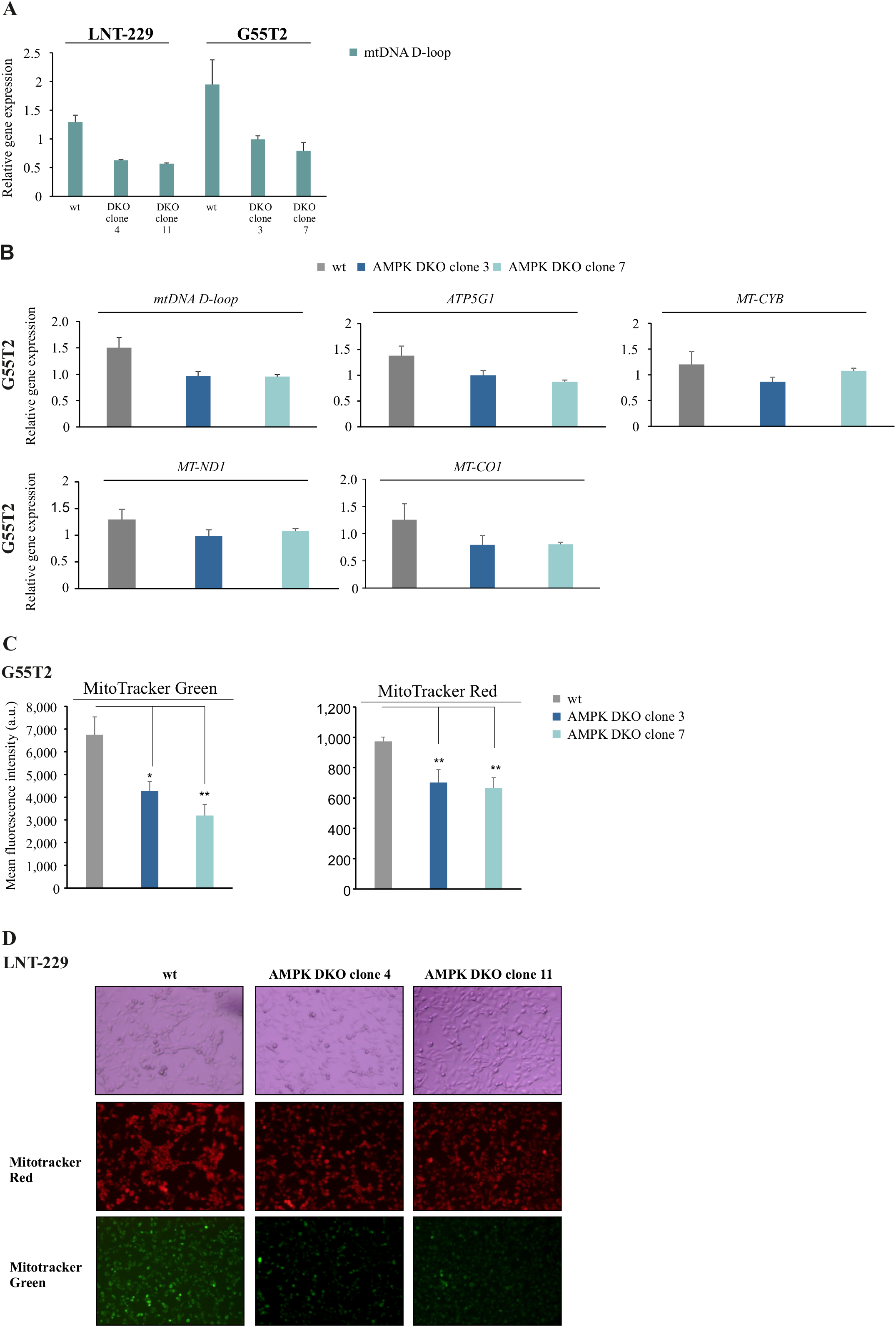
Mitochondrial mass and activity are impaired by AMPK DKO in human GB cell lines. (A) LNT-229 and G55T2 wt and AMPK DKO cells were analysed for the mRNA expression of *mtDNA D-loop* by qPCR. *18S* and *SDHA* were used for normalisation. (B) mRNA expression of mitochondrial encoded as well as mitochondrial associated genes (*mtDNA D-loop*, *ATP5G1*, *MT-CYB*, *MT-ND1* and *MT-CO1*) of G55T2 wt and AMPK DKO cells was determined by qPCR. *18S* and *SDHA* were used as housekeeping genes for normalisation. (C) G55T2 wt and AMPK DKO cells were incubated in serum-free DMEM for 24h. Cells were stained with 100 nM MitoTracker Green or 100 nM MitoTracker Red for 20 min and analysed by flow cytometry. Mean fluorescence intensities are shown (n=3, mean ± SD, *p<0.05, **p<0.01, Student’s t-test). (D) LNT-229 wt and AMPK DKO cells were treated as described in (C). Bright-field (upper panel) and fluorescence microscopy (RFP channel: middle panel, GFP channel: lower panel) were used for analysis (48x magnification). Representative images are shown.

Pharmacological inhibition of oxidative phosphorylation was induced by IACS-010759, which selectively interferes with mitochondrial respiratory complex I ^36–38^. IACS-010759 treatment of LNT-229 wildtype cells led to increased phosphorylation of AMPK and ACC under glucose deprived conditions in normoxia and hypoxia (Supplementary Fig. 2A). Under hypoxic conditions P-AMPK and P-ACC signal intensities were reduced compared to normoxic conditions. Furthermore, oxygen consumption was reduced by IACS-010759 treatment in LNT-229 wildtype and AMPK DKO cells. Moreover, AMPK DKO cells showed reduced oxygen consumption compared to wildtype cells, which was more pronounced under oxidative phosphorylation inhibition (Supplementary Fig. 2B).

### 2-Deoxyglucose treatment increases cell death in AMPK DKO cells

The glycolysis inhibitor 2-deoxyglucose (2-DG) impairs glucose metabolism by accumulation of the unconvertable intermediate 2-deoxyglucose-6-phosphate, which inhibits phosphoglucose isomerase ^39^. We investigated the effects of glycolysis inhibition on AMPK by using AMPK DKO cells. While 2-DG induced AMPK and ACC phosphorylation in LNT-229 wildtype cells in normoxia and hypoxia, no effect on AMPK and ACC phosphorylation was observed in AMPK DKO cells (Fig. 5A). Cell growth analyses showed that 2-DG treatment led to reduced cell densities in wildtype LNT-229 cells but this effect was more pronounced in AMPK DKO cells in both tested glucose/2-DG ratios (Fig. 5B). Under normoxic conditions 2-DG treatment resulted in protection from glucose-starvation induced cell death in wildtype, while the opposite effect was observed in AMPK DKO cells. 2-DG treatment under concomitant glucose and oxygen starvation led to protection from hypoxia-induced cell death in LNT-229 wildtype cells, whereas AMPK DKO cells did not benefit from 2-DG treatment (Fig. 5C).

**Fig. 5:**
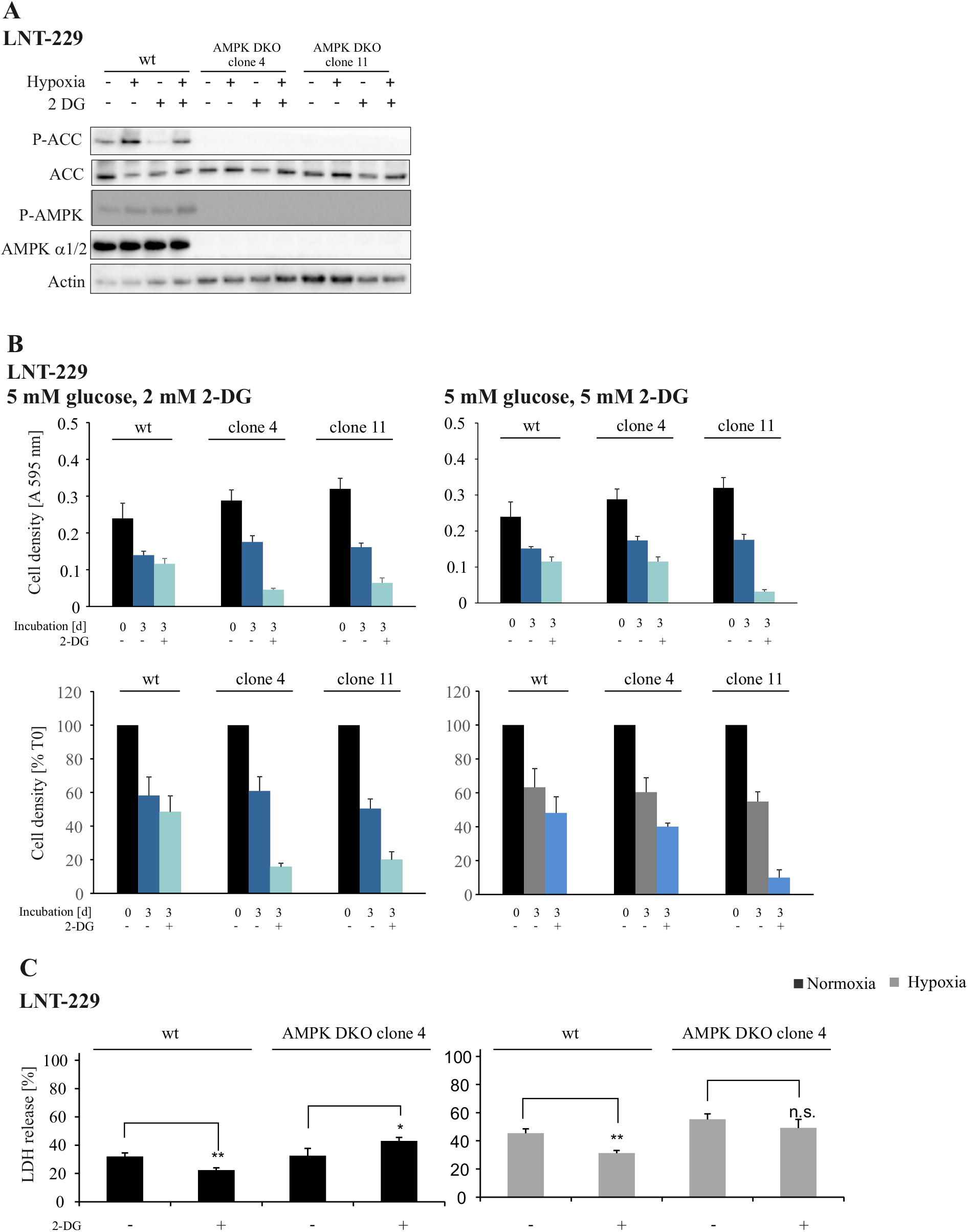
2-DG reduces cell growth in AMPK DKO cells. (A) LNT-229 wildtype and AMPK DKO cells were treated with 5 mM 2-DG in serum-free DMEM containing 2 mM glucose in normoxia or hypoxia (0.1 % O_2_) for 8 h as indicated. Immunoblots were analysed with antibodies for P-ACC, ACC, P-AMPK, AMPK α1/2 and actin. (B) CV staining of LNT-229 wildtype and AMPK DKO cells was performed after treatment with vehicle (DMSO), 2 mM (left panel) or 5 mM (right panel) 2-DG in serum-free medium supplemented with 5 mM glucose for 72 h. Results are shown as absorption at 595 nm and relative to T0, reflecting the respective cell density at the beginning of the experiment. (C) LNT-229 wildtype and AMPK DKO cells were treated with vehicle (DMSO) or 2 mM 2-DG in serum-free DMEM with 2 mM glucose in normoxia or hypoxia (0.1 % O_2_). LDH release assay was used for cell death analysis (n=4, mean ± SD, n.s. not significant, *p<0.05, **p<0.01, Student’s t-test).

### Knockout of both catalytic AMPK subunits impairs tumour formation and leads to prolonged survival in an orthotopic glioma mouse model

The effects of AMPK DKO on tumour formation and tumour progression were investigated in *in vivo* experiments. Chorioallantoic membrane (CAM) assays represent a simple *in vivo* approach to analyse biological effects of tumour formation and progression ^40–42^. The G55T2 orthotopic model has been shown to cause necrotic tumour growth similar to human GB ^43^. Under such conditions, AMPK-mediated effects are presumably more pronounced than in less aggressive tumour models. G55T2 wildtype and AMPK DKO cells were first allowed to grow on the CAM of fertilized chicken eggs (Supplementary Fig. 3A). After 8 days of incubation G55T2 wildtype cells formed significantly larger tumours with up to 70 % increased tumour weights compared to G55T2 AMPK DKO cells (Supplementary Fig. 3B). IHC analysis of FFPE tissue of CAM tumours showed P-AMPK positive staining only in a small fraction of G55T2 AMPK DKO cells and no signal for P-ACC staining (Supplementary Fig. 3C). P-ACC expression was found to be dependent on tumour cell localisation with lower expression in the outer and higher expression in the tumour centre of G55T2 wildtype CAM tumours (Supplementary Fig. 3C). CA IX is known as surrogate marker for hypoxia and its expression is therefore increased in tumour regions with low oxygen supply ^44,45^. IHC staining showed that CA IX expression was induced in G55T2 wildtype CAM tumours, especially in the tumour centre whereas G55T2 AMPK DKO CAM tumours showed only low expression of CA IX (Supplementary Fig. 3D).

In addition to the CAM assays, tumour formation, growth and overall survival was analysed in an orthotopic mouse model. Here, G55T2 AMPK DKO cells showed delayed tumour formation compared to G55T2 wildtype cells in MRI measurements (Fig. 6A). While all animals injected with G55T2 wildtype cells developed tumours that were detectable by MRI at day 18, this was only the case for two of nine mice injected with G55T2 AMPK DKO cells. In line with these results, volumetric analyses based on MRI measurements showed significantly larger tumours in mice injected with G55T2 wildtype cells at day 18 after tumour cell injection (Fig. 6B). With 31 versus 21.5 days in G55T2 AMPK DKO and wildtype tumour bearing mice, survival of mice with G55T2 AMPK DKO tumours was approximately 50% increased (Fig. 6C).

**Fig. 6:**
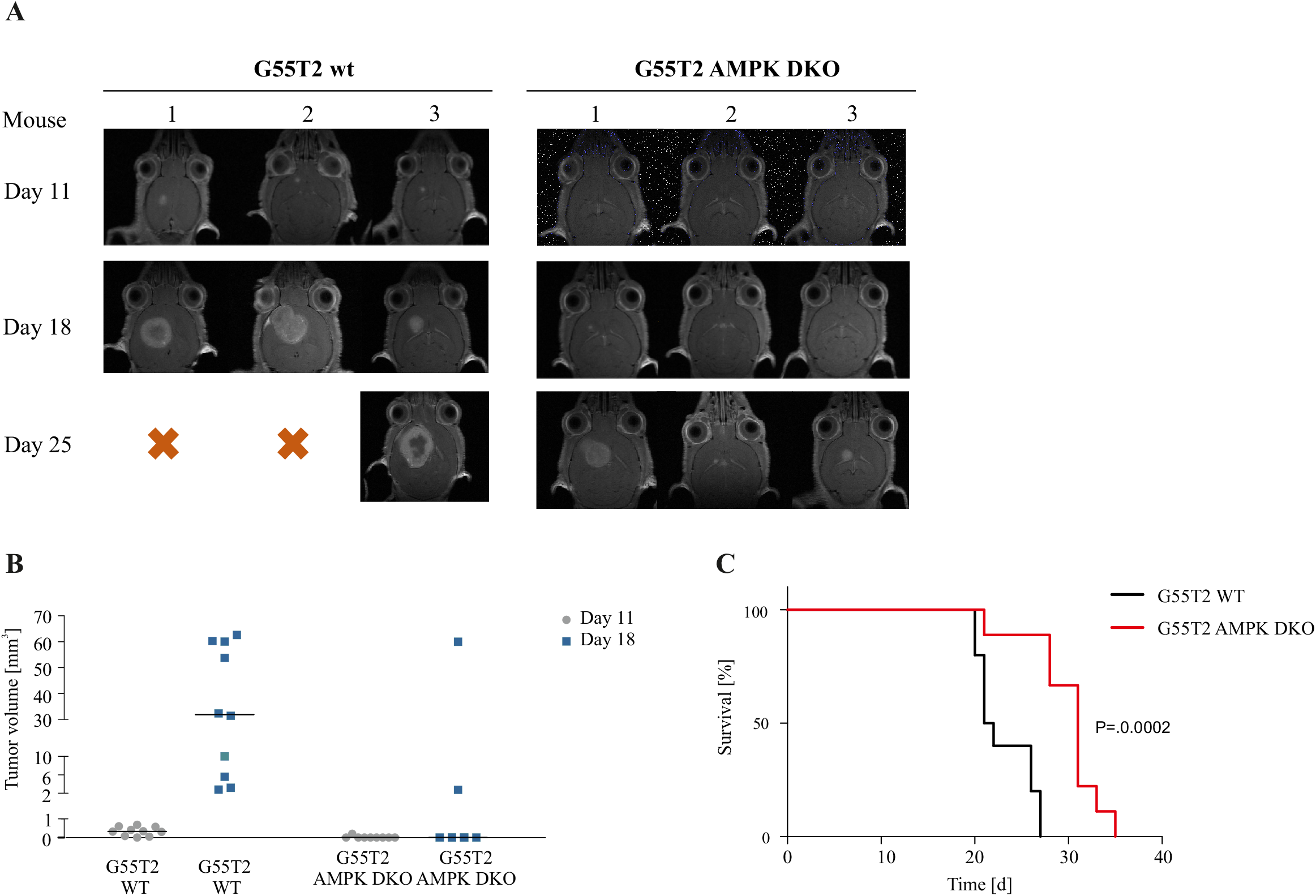
AMPK DKO impairs tumour formation *in vivo*. (A) Athymic nude mice were intracranially injected with 1 x 10^5^ G55T2 wildtype or AMPK DKO cells (n=10 mice per group). MRI measurements were performed on day 11, 18 and 25 after tumour cell injection. Images of 3 mice per group are shown exemplarily. (B) Tumour volumes were calculated based on MRI measurements on day 11 and 18 using the ITK Snap software. (C) Survival of mice injected with G55T2 wildtype or AMPK DKO cells was analysed by Kaplan-Meier Plot. Significance was tested using Log-Rank test.

## Discussion

For sustained tumour cell survival and proliferation within the GB microenvironment, precise sensing of the cellular energy state is an important prerequisite to coordinate metabolism. AMPK orchestrates metabolism by increasing catabolism and inhibiting anabolism ^11,46,47^ with potentially specific relevance under conditions of the GB microenvironment. Accordingly, chronic AMPK activation has been reported in GB cells ^18^ and therefore AMPK could represent a promising target for anti-glioma therapies.

With this study, we provide evidence that functional AMPK is essential for adaptation of human GB cells to energy starvation conditions with direct effects on GB cell survival (Fig. 7). AMPK was induced under nutrient-deprived as well as hypoxic conditions (Fig. 1B), but double knockout of the catalytic subunits α1 and α2 sensitised human GB cells to cell death induced by glucose starvation-mediated and hypoxia-induced cell death, both mimicking realistic conditions of the GB microenvironment (Fig. 2). These results confirm previous studies which demonstrated the significance of AMPK in GB cells and showed increased levels of the AMPK subunits α1, β1 and γ1 on mRNA level compared to normal brain tissue as well as increased levels of phosphorylated AMPK in GB cells ^18^. Gene suppression of the β1 regulatory AMPK subunit has been shown to result in downregulation of cellular bioenergetics and reduced tumour growth *in vivo* ^18^. Nevertheless, in this model, phosphorylation of AMPK was not fully inhibited because of a sustained catalytic AMPK activity. Knockout of the catalytic subunits α1 and α2 therefore could more precisely reveal AMPK specific effects. Transfection of AMPK DKO cells with *PRKAA2*, encoding for the α2 subunit, partially rescued cell death under glucose starvation and hypoxia (Fig. 3C, D). Of note, transfection with PRKAA2 did not lead to protein levels comparable to the wildtype AMPK protein level and thus could explain the partial rescue of those cells (Fig. 3A). However, compared to wildtype GB cells, it was also evident that both catalytic subunits are required for complete AMPK activity and adaptation to energy stress conditions.

**Fig. 7:**
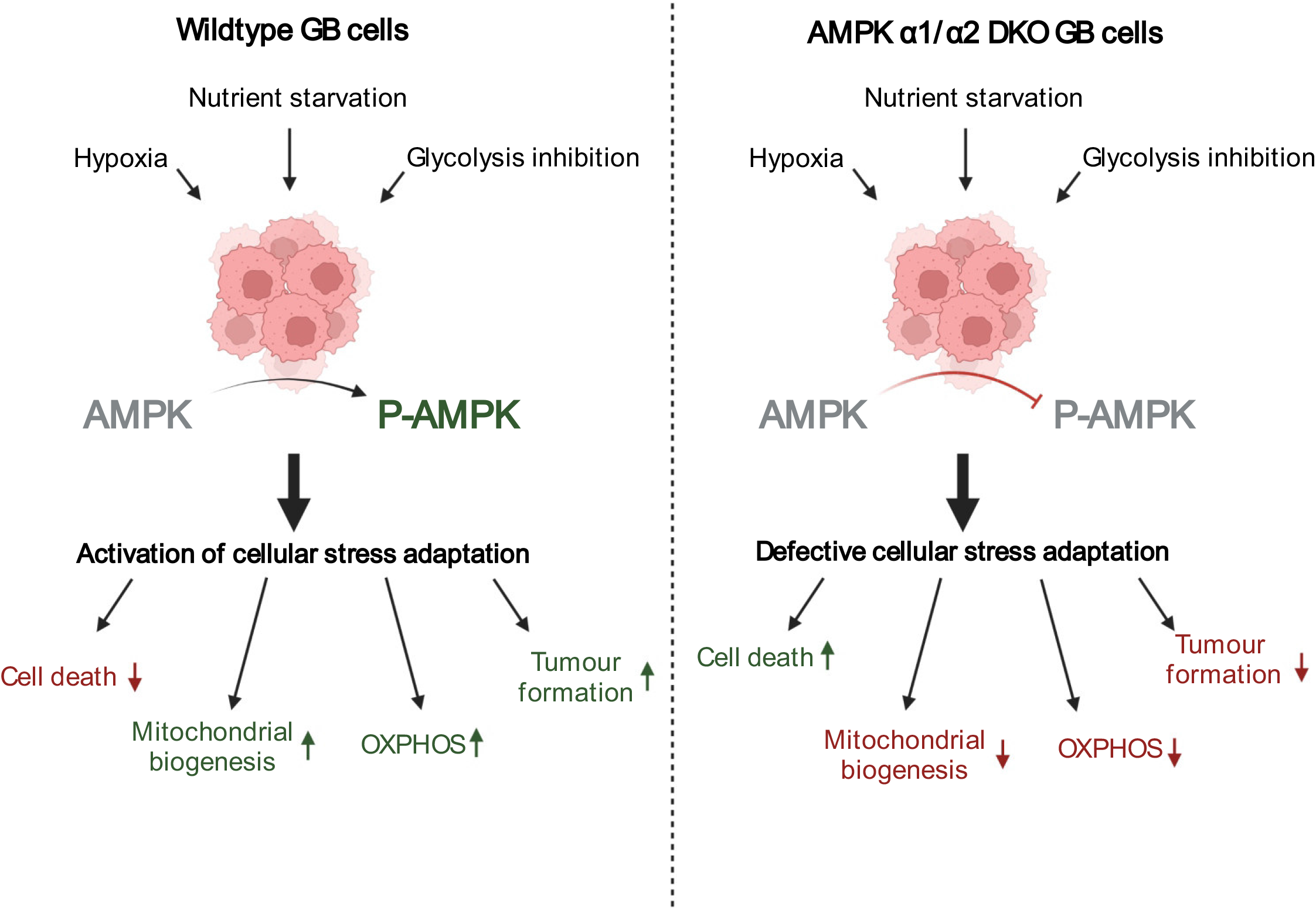
Double knockout of the AMPK catalytic subunits renders GB cells more sensitive to conditions of the tumour microenvironment. GB cells are challenged by the surrounding tumour microenvironment (low oxygen and nutrient availability) and therefore activate AMPK to overcome energy starvation conditions (left panel). By activation of AMPK-dependent cellular adaptation programs, catabolic processes (OXPHOS) and mitochondrial biogenesis are increased resulting in an increased tumour cell survival which promotes tumour formation. Double knockout of AMPK catalytic subunits α1 and α2 impairs AMPK signalling, leading to an absent response to energy starvation conditions (right panel). In consequence, tumour cell metabolism is no longer able to adapt to nutrient- and oxygen-deprived conditions which is linked to an increased GB cell death and impaired tumour formation. Figure created with biorender.com.

We also found similar effects in the primary GB cell setting by employing pharmacological AMPK inhibition with BAY3827 (Fig. 2C, D, Supplementary Fig. 1A-C). In contrast to the frequently used AMPK inhibitor Compound C, which is known to lack AMPK specificity, BAY3827 is a selective and potent inhibitor of AMPK subunit PRKAA1 with low potency off-target activity for RPS6K1 ^34^. In both, LNT-229 and primary GB cells (P3NS), BAY3827 treatment increased cell death under glucose starvation and hypoxia as it was also observed in the AMPK DKO model (Fig. 2C, D, Supplementary Fig. 1A-C). Moreover, BAY3827 treatment did not influence hypoxia-induced cell death rates in AMPK DKO cells arguing for low off-target effects (Supplementary Fig. 1C). Selective pharmacological AMPK inhibition could therefore be beneficial for GB treatment by rendering cells more susceptible towards conditions of the tumour microenvironment.

AMPK modulates mitochondrial biogenesis and homeostasis dependent on the cellular energy state ^35^. Only little is known about the role of AMPK for mitochondria in GB cells. In the β1 gene suppression model, primary GB cells were characterized by reduced mitochondrial activity and mass ^18^. Here, we demonstrate that inhibition of catalytic AMPK activity induced by α1/α2 knockdown led to significant reduction of mitochondrial DNA content and mass (Fig. 4A, C MitoTracker Green, D) together with reduced mitochondrial membrane potential indicating lower mitochondrial activity (Fig. 4C MitoTracker Red, D). In line with this observation, mRNA levels of mitochondrial encoded and mitochondria associated genes decreased in AMPK DKO cells compared to wildtype GB cells (Fig. 4B). In this context, PGC-1α, encoded by *PPARGC1A*, as master regulator of mitochondrial biogenesis has been found to be AMPK-dependently activated *via* p38MAPK activation leading to an enhanced mitochondrial activity and biogenesis to improve ATP synthesis to overcome energy depletion under nutrient-deprived conditions ^15^. AMPK-dependent regulation of mitochondrial encoded genes in GB cells with reduced mitochondrial mass and activity accompanied by a reduced oxygen consumption found in AMPK DKO cells (Supplementary Fig. 2B). The essential role of AMPK to sustain mitochondrial activity for ATP production under energy starvation conditions, was confirmed by the fact that OXPHOS inhibition and AMPK DKO both resulted in a comparable reduction of oxygen consumption in LNT-229 cells (Supplementary Fig. 2B). Taken together, deregulated mitochondrial capacity to produce ATP appears to be an important consequence of defective AMPK activity.

Aerobic glycolysis is known to be an important route to metabolise glucose in tumour cells ^48,49^. Therefore, tumour cells are more susceptible to the treatment with glycolysis inhibitors like 2-DG ^50^. In consequence of glycolysis inhibition, ATP is depleted and AMPK is activated ^8,51^. Our results demonstrate that AMPK DKO cells show an impaired adaptation to energy starvation conditions and thus it seems plausible that these cells are more sensitive to glycolysis inhibition than wildtype cells. In line, LNT-229 AMPK DKO cells showed decreased cell growth under glucose-deprived conditions combined with 2-DG treatment compared to LNT-229 wildtype cells (Fig. 5B). Moreover, treatment of AMPK DKO cells with 2-DG led to increased cell death compared to LNT-229 wildtype cells, which were protected from glucose starvation-mediated cell death because of an increased AMPK activation resulting in the inhibition of catabolic processes and activation of ATP generation to restore cellular energy storage. AMPK inhibition together with glycolysis inhibition (e.g. with 2-DG) could therefore be beneficial to render GB cells more susceptible to conditions of the tumour microenvironment and to increase cell death under treatment conditions.

Findings of our *in vivo* models emphasise the role of AMPK for tumour formation. In line with previous studies, where gene suppression of the regulatory β1 subunit was sufficient to induce reduced survival in mice, we found that knockout of the catalytic AMPK subunits led to impaired tumour formation of CAM tumours and in mouse experiments (Fig. 6A, B). Moreover, survival of AMPK DKO tumour bearing mice was significantly prolonged (Fig. 6C). Together these results indicate that AMPK is essential especially for tumour progression. In this context, AMPK could be essential for adaptation to conditions of the tumour microenvironment characterised by low nutrient and oxygen supply. Further experiments should investigate the role of AMPK for tumour growth *in vivo* in more detail and address the question whether concomitant AMPK inhibition and glycolysis inhibition could increase cell death of GB cells. In summary, adaptation to the tumour microenvironment is a central prerequisite for sustained tumour growth in solid cancers. Our study demonstrates that AMPK mediates adaptation of GB cells to energy stress conditions. Therefore, AMPK inhibition could be a promising strategy for GB treatment either as monotherapy or to sensitise tumour cells to other metabolically active therapies.

## Declarations

## Acknowledgements

The DCP compounds BAY974 and BAY3827 were supplied by the Structural Genomics Consortium under an Open Science Trust Agreement: http://www.thesgc.org/click-trust. We thank Simone Niclou and Anna Golebiewska for contributing primary cells. The authors thank Niklas Lohfink for the technical support.

## Availability of data and materials

The datasets used or analysed during the current study are available from the corresponding author on reasonable request.

## Authors’ contributions

D.S, S.N., M.M., L.S., P.N.H., C.M., D.N., M.W.R. and J.P.S. conceived the study and designed the experiments. N.I.L., B.S., P.S.Z., M.I.S., A.-L.L., T.A. and M.W.R., performed the experiments. N.I.L., B.S., P.S.Z., M.I.S., A.-L.L. and M.W.R. analysed the data. N.I.L. and M.W.R. wrote the manuscript. All authors helped to draft the manuscript and read and approved the final version.

## References

1. Monteiro AR, Hill R, Pilkington GJ, Madureira PA. The Role of Hypoxia in Glioblastoma Invasion. Cells. 2017;6(4). doi:10.3390/cells6040045

2. Stupp R, Mason WP, van den Bent MJ, et al. Radiotherapy plus concomitant and adjuvant temozolomide for glioblastoma. N Engl J Med. 2005;352(10):987–996. doi:10.1056/NEJMoa043330

3. Hanahan D, Weinberg RA. Hallmarks of cancer: the next generation. Cell. 2011;144(5):646–674. doi:10.1016/j.cell.2011.02.013

4. Weyandt JD, Thompson CB, Giaccia AJ, Rathmell WK. Metabolic Alterations in Cancer and Their Potential as Therapeutic Targets. Am Soc Clin Oncol Educ Book. 2017;37:825–832. doi:10.14694/EDBK_175561

5. Hardie DG, Schaffer BE, Brunet A. AMPK: An Energy-Sensing Pathway with Multiple Inputs and Outputs. Trends Cell Biol. 2016;26(3):190–201. doi:10.1016/j.tcb.2015.10.013

6. Lin S-C, Hardie DG. AMPK: Sensing Glucose as well as Cellular Energy Status. Cell Metab. 2018;27(2):299–313. doi:10.1016/j.cmet.2017.10.009

7. Zhang C-S, Hawley SA, Zong Y, et al. Fructose-1,6-bisphosphate and aldolase mediate glucose sensing by AMPK. Nature. 2017;548:112 EP–. doi:10.1038/nature23275

8. Garcia D, Shaw RJ. AMPK: Mechanisms of Cellular Energy Sensing and Restoration of Metabolic Balance. Mol Cell. 2017;66(6):789–800. doi:10.1016/j.molcel.2017.05.032

9. Xiao B, Sanders MJ, Underwood E, et al. Structure of mammalian AMPK and its regulation by ADP. Nature. 2011;472(7342):230–233. doi:10.1038/nature09932

10. Hardie DG, Ross FA, Hawley SA. AMPK: a nutrient and energy sensor that maintains energy homeostasis. Nat Rev Mol Cell Biol. 2012;13(4):251–262. doi:10.1038/nrm3311

11. Hardie DG, Pan DA. Regulation of fatty acid synthesis and oxidation by the AMP-activated protein kinase. Biochem Soc Trans. 2002;30(Pt 6):1064–1070.

12. Laplante M, Sabatini DM. mTOR signaling in growth control and disease. Cell. 2012;149(2):274–293. doi:10.1016/j.cell.2012.03.017

13. Herzig S, Shaw RJ. AMPK: guardian of metabolism and mitochondrial homeostasis. Nat Rev Mol Cell Biol. 2018;19(2):121–135. doi:10.1038/nrm.2017.95

14. Zhang C-S, Lin S-C. AMPK Promotes Autophagy by Facilitating Mitochondrial Fission. Cell Metab. 2016;23(3):399–401. doi:10.1016/j.cmet.2016.02.017

15. Chaube B, Malvi P, Singh SV, Mohammad N, Viollet B, Bhat MK. AMPK maintains energy homeostasis and survival in cancer cells via regulating p38/PGC-1α-mediated mitochondrial biogenesis. Cell Death Discov. 2015;1:15063–. doi:10.1038/cddiscovery.2015.63

16. Faubert B, Vincent EE, Poffenberger MC, Jones RG. The AMP-activated protein kinase (AMPK) and cancer: many faces of a metabolic regulator. Cancer Lett. 2015;356(2 Pt A):165–170. doi:10.1016/j.canlet.2014.01.018

17. Jiang H, Liu W, Zhan S-K, et al. GSK621 Targets Glioma Cells via Activating AMP-Activated Protein Kinase Signalings. PLoS ONE. 2016;11(8):e0161017. doi:10.1371/journal.pone.0161017

18. Chhipa RR, Fan Q, Anderson J, et al. AMP kinase promotes glioblastoma bioenergetics and tumour growth. Nat Cell Biol. 2018;20(7):823–835. doi:10.1038/s41556-018-0126-z

19. Shackelford DB, Abt E, Gerken L, et al. LKB1 inactivation dictates therapeutic response of non-small cell lung cancer to the metabolism drug phenformin. Cancer Cell. 2013;23(2):143–158. doi:10.1016/j.ccr.2012.12.008

20. Bain J, Plater L, Elliott M, et al. The selectivity of protein kinase inhibitors: a further update. Biochem J. 2007;408(3):297–315. doi:10.1042/BJ20070797

21. Dasgupta B, Seibel W. Compound C/Dorsomorphin: Its Use and Misuse as an AMPK Inhibitor. Methods Mol Biol. 2018;1732:195–202. doi:10.1007/978-1-4939-7598-3_12

22. Liu X, Chhipa RR, Nakano I, Dasgupta B. The AMPK inhibitor compound C is a potent AMPK-independent antiglioma agent. Mol Cancer Ther. 2014;13(3):596–605. doi:10.1158/1535-7163.MCT-13-0579

23. Ronellenfitsch MW, Brucker DP, Burger MC, et al. Antagonism of the mammalian target of rapamycin selectively mediates metabolic effects of epidermal growth factor receptor inhibition and protects human malignant glioma cells from hypoxia-induced cell death. Brain. 2009;132(Pt 6):1509–1522. doi:10.1093/brain/awp093

24. Eckerich C, Schulte A, Martens T, Zapf S, Westphal M, Lamszus K. RON receptor tyrosine kinase in human gliomas: expression, function, and identification of a novel soluble splice variant. J Neurochem. 2009;109(4):969–980. doi:10.1111/j.1471-4159.2009.06027.x

25. Wischhusen J, Naumann U, Ohgaki H, Rastinejad F, Weller M. CP-31398, a novel p53-stabilizing agent, induces p53-dependent and p53-independent glioma cell death. Oncogene. 2003;22(51):8233–8245. doi:10.1038/sj.onc.1207198

26. Lorenz NI, Sittig ACM, Urban H, et al. Activating transcription factor 4 mediates adaptation of human glioblastoma cells to hypoxia and temozolomide. Sci Rep. 2021;11(1):14161. doi:10.1038/s41598-021-93663-1

27. SGC Frankfurt. http://www.thesgc.org/click-trust

28. Golebiewska A, Hau A-C, Oudin A, et al. Patient-derived organoids and orthotopic xenografts of primary and recurrent gliomas represent relevant patient avatars for precision oncology. Acta Neuropathol. 2020;140(6):919–949. doi:10.1007/s00401-020-02226-7

29. Wanka C, Brucker DP, Bähr O, et al. Synthesis of cytochrome C oxidase 2: a p53-dependent metabolic regulator that promotes respiratory function and protects glioma and colon cancer cells from hypoxia-induced cell death. Oncogene. 2012;31(33):3764–3776. doi:10.1038/onc.2011.530

30. Grady JE, Lummis WL, Smith CG. An improved tissue culture assay. III. Alternate methods for measuring cell growth. Cancer Res. 1960;20:1114–1117.

31. Steinbach JP, Eisenmann C, Klumpp A, Weller M. Co-inhibition of epidermal growth factor receptor and type 1 insulin-like growth factor receptor synergistically sensitizes human malignant glioma cells to CD95L-induced apoptosis. Biochem Biophys Res Commun. 2004;321(3):524–530. doi:10.1016/j.bbrc.2004.06.175

32. Thiepold A-L, Lorenz NI, Foltyn M, et al. Mammalian target of rapamycin complex 1 activation sensitizes human glioma cells to hypoxia-induced cell death. Brain. 2017;140(10):2623–2638. doi:10.1093/brain/awx196

33. Yushkevich PA, Piven J, Hazlett HC, et al. User-guided 3D active contour segmentation of anatomical structures: significantly improved efficiency and reliability. Neuroimage. 2006;31(3):1116–1128. doi:10.1016/j.neuroimage.2006.01.015

34. Lemos C, Schulze VK, Baumgart SJ, et al. The potent AMPK inhibitor BAY-3827 shows strong efficacy in androgen-dependent prostate cancer models. Cell Oncol (Dordr). 2021;44(3):581–594. doi:10.1007/s13402-020-00584-8

35. Toyama EQ, Herzig S, Courchet J, et al. AMP-activated protein kinase mediates mitochondrial fission in response to energy stress. Science. 2016;351(6270):275–281. doi:10.1126/science.aab4138

36. Tsuji A, Akao T, Masuya T, Murai M, Miyoshi H. IACS-010759, a potent inhibitor of glycolysis-deficient hypoxic tumor cells, inhibits mitochondrial respiratory complex I through a unique mechanism. J Biol Chem. 2020;295(21):7481–7491. doi:10.1074/jbc.RA120.013366

37. Molina JR, Sun Y, Protopopova M, et al. An inhibitor of oxidative phosphorylation exploits cancer vulnerability. Nat Med. 2018;24(7):1036–1046. doi:10.1038/s41591-018-0052-4

38. Vangapandu HV, Alston B, Morse J, et al. Biological and metabolic effects of IACS-010759, an OxPhos inhibitor, on chronic lymphocytic leukemia cells. Oncotarget. 2018;9(38):24980–24991. doi:10.18632/oncotarget.25166

39. Pajak B, Siwiak E, Sołtyka M, et al. 2-Deoxy-d-Glucose and Its Analogs: From Diagnostic to Therapeutic Agents. Int J Mol Sci. 2019;21(1). doi:10.3390/ijms21010234

40. Lokman NA, Elder ASF, Ricciardelli C, Oehler MK. Chick Chorioallantoic Membrane (CAM) Assay as an In Vivo Model to Study the Effect of Newly Identified Molecules on Ovarian Cancer Invasion and Metastasis. Int J Mol Sci. 2012;13(8):9959–9970. doi:10.3390/ijms13089959

41. Richardson M, Singh G. Observations on the use of the avian chorioallantoic membrane (CAM) model in investigations into angiogenesis. Curr Drug Targets Cardiovasc Haematol Disord. 2003;3(2):155–185. doi:10.2174/1568006033481492

42. Deryugina EI, Quigley JP. Chick embryo chorioallantoic membrane model systems to study and visualize human tumor cell metastasis. Histochem Cell Biol. 2008;130(6):1119–1130. doi:10.1007/s00418-008-0536-2

43. Seidel S, Garvalov BK, Wirta V, et al. A hypoxic niche regulates glioblastoma stem cells through hypoxia inducible factor 2 alpha. Brain. 2010;133(Pt 4):983–995. doi:10.1093/brain/awq042

44. Vaupel P, Mayer A. Hypoxia in cancer: significance and impact on clinical outcome. Cancer Metastasis Rev. 2007;26(2):225–239. doi:10.1007/s10555-007-9055-1

45. McLendon R. Carbonic anhydrase IX as a marker of hypoxia in gliomas: A narrative review. Glioma. 2020;3(3):97. doi:10.4103/glioma.glioma_19_20

46. Jeon S-M, Chandel NS, Hay N. AMPK regulates NADPH homeostasis to promote tumour cell survival during energy stress. Nature. 2012;485(7400):661–665. doi:10.1038/nature11066

47. Kim J, Kundu M, Viollet B, Guan K-L. AMPK and mTOR regulate autophagy through direct phosphorylation of Ulk1. Nat Cell Biol. 2011;13(2):132–141. doi:10.1038/ncb2152

48. Warburg O. On respiratory impairment in cancer cells. Science. 1956;124(3215):269–270.

49. Vander Heiden MG, Cantley LC, Thompson CB. Understanding the Warburg effect: the metabolic requirements of cell proliferation. Science. 2009;324(5930):1029–1033. doi:10.1126/science.1160809

50. Aft RL, Zhang FW, Gius D. Evaluation of 2-deoxy-D-glucose as a chemotherapeutic agent: mechanism of cell death. Br J Cancer. 2002;87(7):805–812. doi:10.1038/sj.bjc.6600547

51. Barialai L, Strecker MI, Luger A-L, et al. AMPK activation protects astrocytes from hypoxia-induced cell death. Int J Mol Med. 2020;45(5):1385–1396. doi:10.3892/ijmm.2020.4528

