## Supplementary Information for "AMP-kinase mediates adaptation of glioblastoma cells to conditions of the tumour microenvironment"

#### Supplementary Methods:

##### Chorioallantoic membrane (CAM) Assay

Fertilized chicken eggs (LSL Rhein-Main, Dieburg, Germany) were incubated at 37°C for seven days after fertilization. 2 x $10^6$ cells per egg were diluted in 10 µl DMEM and 10 µl Corning Matrigel (Corning, Amsterdam, Netherlands) and placed onto a blood vessel of the chorioallantoic membrane. Tumours were isolated after further seven days of incubation. Tumour weights were documented and tumours were further analysed by immunohistological staining.

##### Measurement of oxygen consumption

Cells were seeded and incubated overnight to ensure cell attachment. Cultured cells treated as indicated and were overlaid with sterile paraffin oil. Oxygen consumption was determined with a fluorescence-based assay (PreSens, Regensburg, Germany) [1].

##### Immunohistological staining

Immunohistochemical (IHC) analyses were performed with formalin-fixed paraffin-embedded (FFPE) tissue of CAM tumour models. The following antibodies were used: anti-CA9 (#5649, clone D47G3, dilution 1:100, Cell Signaling Technology, Danvers, MA, USA), anti-p-ACC (Ser79) (#3661, dilution 1:200, Cell Signaling Technology, Danvers, MA, USA) and anti-p-AMPK (Thr172) (#2535, dilution 1:100, Cell Signaling Technology, Danvers, MA, USA). Tissue blocks were cut in slices of 3 µm thickness using a microtome (Leica Microsystems

Nussloch GmbH, Nussloch, Germany) and placed onto SuperFrost slides (Thermo Scientific, Dreieich, Germany). IHC was performed according to standardised protocols using Leica Bond RX automated immunostaining system (Leica Biosystems Nussloch GmbH, Nussloch, Germany). IHC stainings were analysed using a light microscope (BX41, Olympus, Hamburg, Germany).

#### References:


1. Thiepold A-L, Lorenz NI, Foltyn M, et al. Mammalian target of rapamycin complex 1 activation sensitizes human glioma cells to hypoxia-induced cell death. Brain. 2017;140(10):2623-2638. doi:10.1093/brain/awx196


### Supplementary Figure Legends:

#### Supplementary Fig. 1: Pharmacological AMPK inhibition affects cell death under glucose starvation- and hypoxia-induced cell death in human GB cell lines.

(A) LNT-229 cells were treated with vehicle, 1 µM BAY974 (negative control) or 1 µM BAY3827 in serum-free DMEM containing 2 mM glucose in normoxia or hypoxia (0.1 % $O_2$) for 8 h as indicated. Immunoblot analysis was performed with antibodies for P-ACC, P-AMPK, AMPK and actin. (B) LNT-229 cells were treated with vehicle (DMSO), 1 µM BAY974 or 1 µM BAY3827 in glucose-free medium as indicated. PI staining was used for cell death analysis and quantified by FACS measurement (n=3, mean ± SD, **p<0.01, Student's t-test). (C) LNT-229 wildtype and AMPK DKO cells were incubated in serum-free medium containing 2 mM glucose and vehicle (DMSO), 1 µM BAY974 or 1 µM BAY3927 under normoxic or hypoxic (0.1 % $O_2$) conditions as indicated. Cell death was analysed by LDH release assay (n=4, mean ± SD, n.s. not significant, **p<0.01, Student's t-test).

#### Supplementary Fig. 2: Inhibition of oxidative phosphorylation induces AMPK activation in human GB cells.

(A) LNT-229 wildtype and AMPK DKO cells were treated with vehicle (DMSO) or 100 nM IACS-010759 in serum-free medium containing 2 mM glucose in normoxia and hypoxia (0.1 % $O_2$) for 8 h. Immunoblot analysis was performed using antibodies for P-ACC, P-AMPK, AMPK α1/2 and actin. (B) LNT-229 cells were treated with vehicle (DMSO) or 100 nM IACS-010759 in serum-free DMEM without glucose restriction. Oxygen consumption was measured by a fluorescence-based assay.

#### Supplementary Fig. 3: CAM tumours of AMPK DKO cells show reduced tumour weights and impaired metabolic adaptation.

(A, B) Chicken eggs were injected with 2 x $10^6$ G55T2 wt or AMPK DKO cells on day 7 post fertilization and incubated for further 8 days. Tumours were isolated on day 15, weights were measured and tumours were fixed for immunohistochemical analysis. Images of isolated tumours are shown (scale bar: 5 mm). (C) G55T2 wt and AMPK DKO CAM tumours were analysed immunohistochemically with antibodies for P-AMPK and P-ACC. Scale bar represents 200 µm (10x magnification, upper and middle panel) or 1 mm (2x magnification, bottom panel). (D) Isolated CAM tumours were stained for CA IX. Scale bar represents 200 µm (upper panel), 1 mm (2x magnification, bottom panel).

**Supplementary Fig. 1**

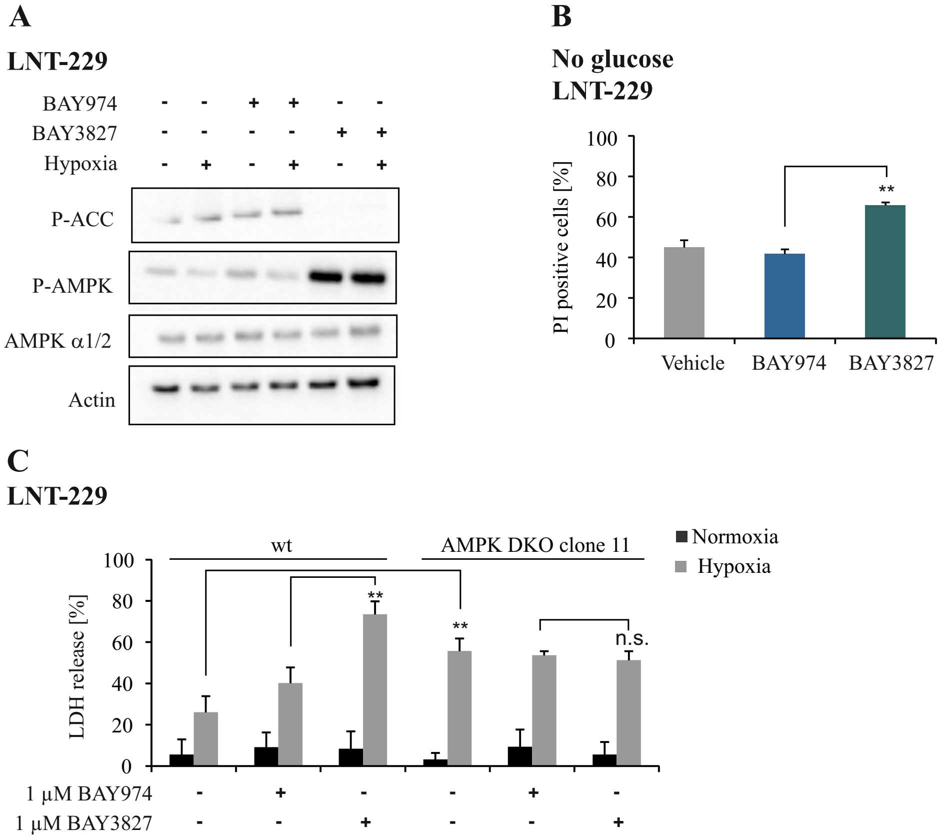

#### Supplementary Fig. 2

**A**

**LNT-229**

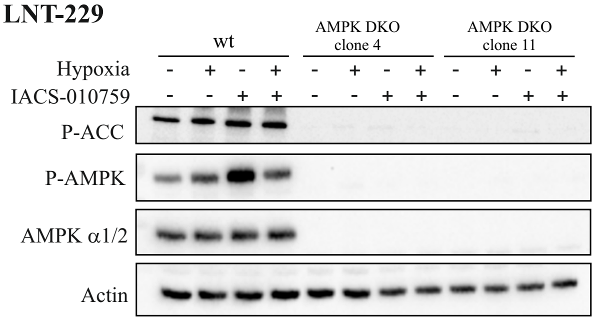


**B**

**LNT-229**

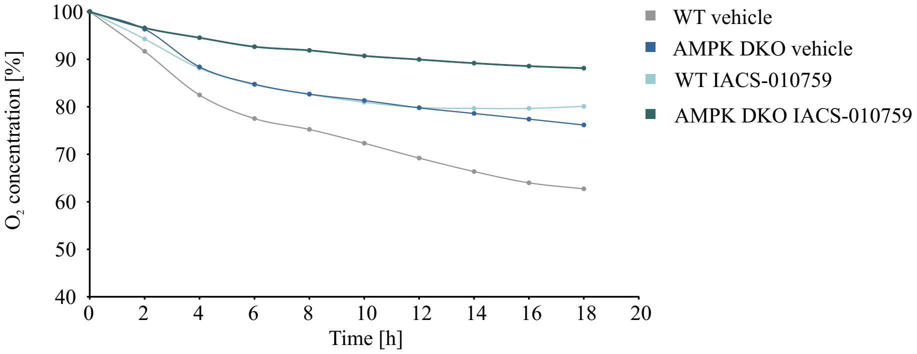

#### Supplementary Fig. 3

### A

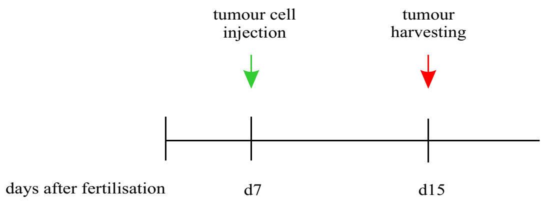


### B

##### G55T2 CAM tumours

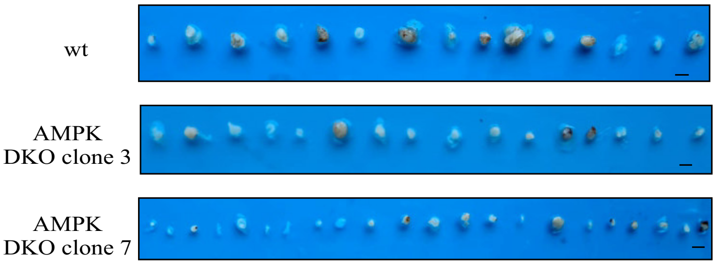


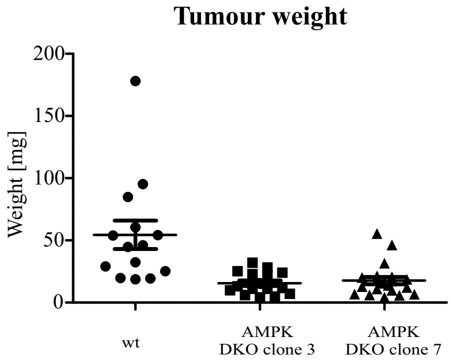


### C

##### G55T2 CAM tumours

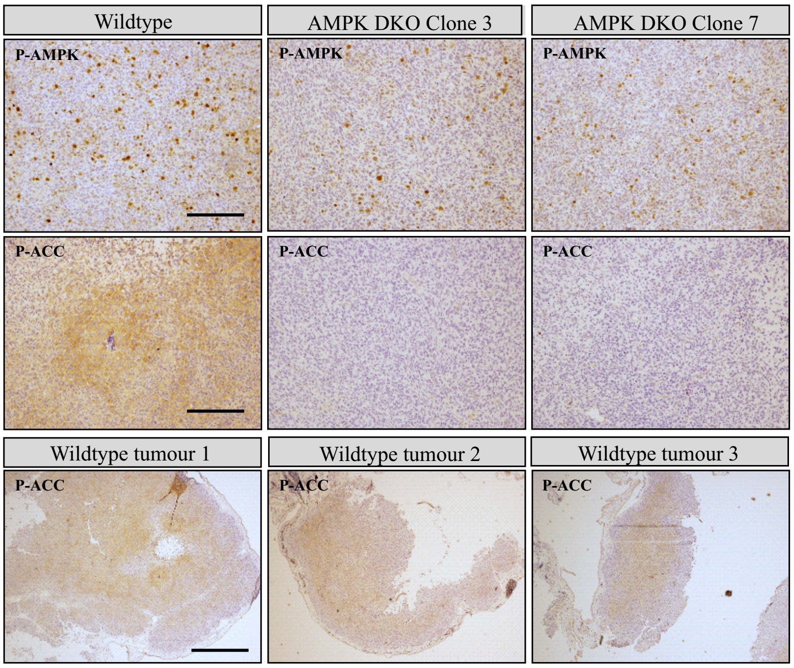


### D

##### G55T2 CAM tumours

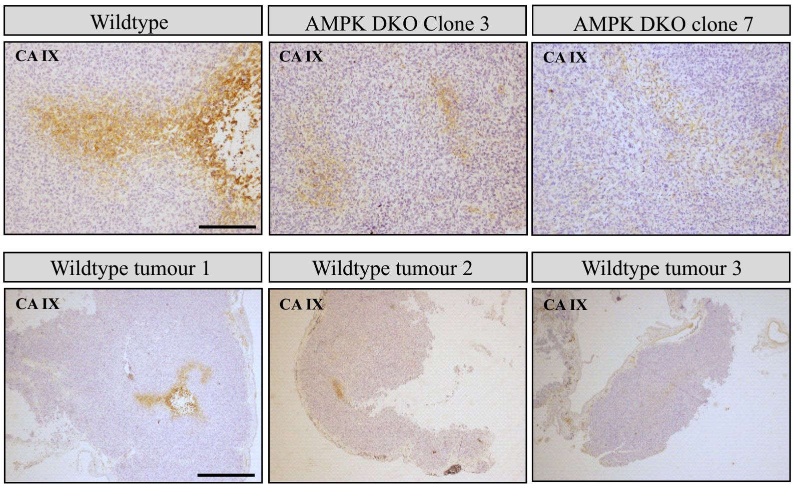
